## Supplementary material for "Understanding the population structure of *Moraxella catarrhalis* using core genome multilocus sequence typing (cgMLST) and a life identification number (LIN) code classification system": SuppFigures

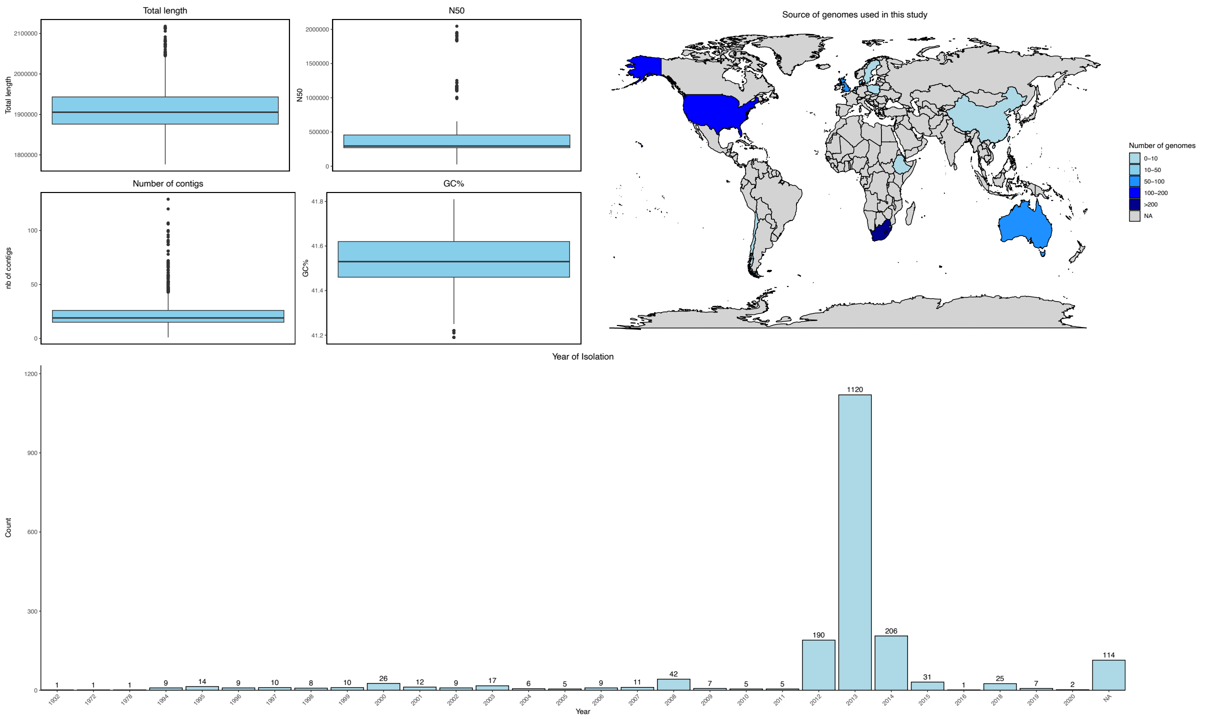


**Supplementary Figure 1.** Characteristics of the 1,913 *M. catarrhalis* genomes that passed quality control and were analysed in this study.

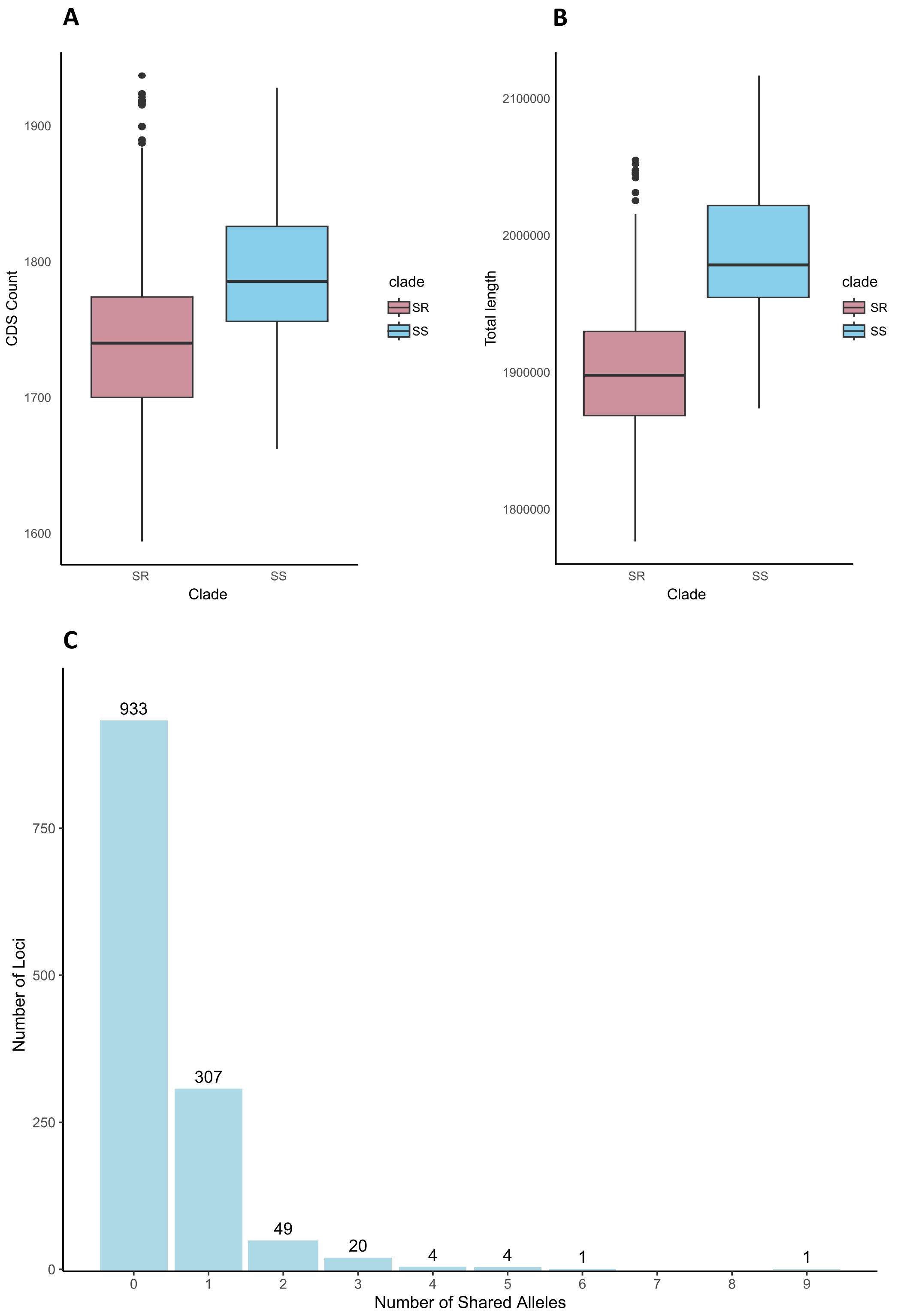


**Supplementary Figure 2**. (A) Box plot showing the distribution of coding sequences in SR and SS genomes. (B) Box plot comparing genome length distributions in SR and SS genomes. (C) Number of shared alleles between SR and SS among the 1,319 core genes. Note: CDS, coding sequence; SR, seroresistant; SS, serosensitive.

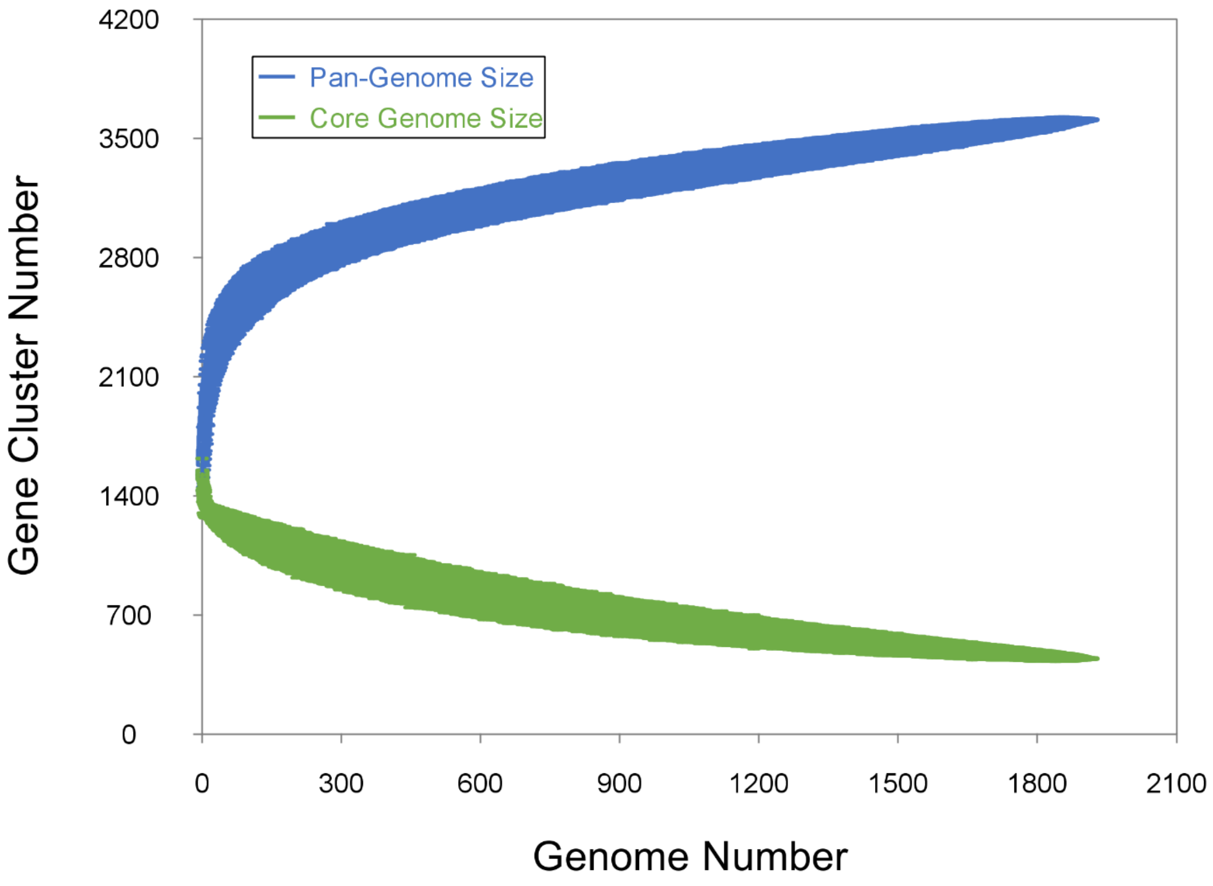


**Supplementary Figure 3**. Rarefaction curves of the pan-genome size and core genome size of *M. catarrhalis,* relative to the number of genomes analysed.

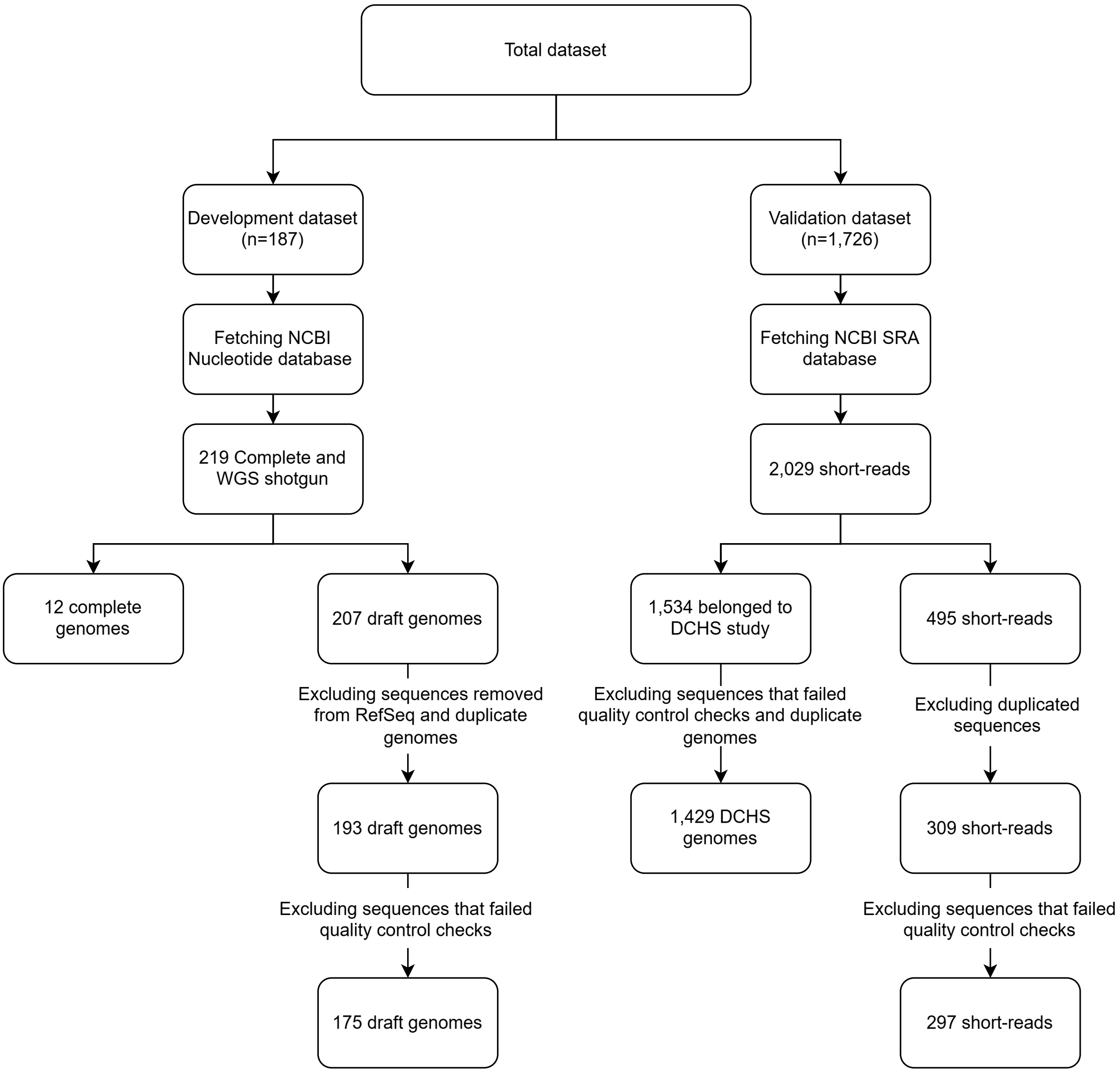


**Supplementary Figure 4.** Schematic overview of the selection and filtering process of *M. catarrhalis* genomes for the development and validation datasets.
